# Phylogeny of the Australian *Solanum dioicum* group using seven nuclear genes: Testing Symon’s fruit and seed dispersal hypotheses

**DOI:** 10.1101/462945

**Authors:** Christopher T. Martine, Ingrid E. Jordon-Thaden, Angela J. McDonnell, Jason T. Cantley, Daniel S. Hayes, Morgan D. Roche, Emma S. Frawley, Ian S. Gilman, David C. Tank

## Abstract

The dioecious and andromonoecious *Solanum* taxa (previously described as the “*S. dioicum* group”) of the Australian Monsoon Tropics have been the subject of phylogenetic and taxonomic study for decades, yet much of their basic biology is still unknown. This is especially true for plant-animal interactions, including the influence of fruit form and calyx morphology on seed dispersal. We combine field/greenhouse observations and specimen-based study with phylogenetic analysis of seven nuclear regions obtained via a microfluidic PCR-based enrichment strategy and high-throughput sequencing, and present the first intron-containing nuclear gene dataset in the genus *Solanum* and the first species-tree hypothesis for the *S. dioicum* group. Our results suggest that epizoochorous trample burr seed dispersal (strongly linked to calyx accrescence) is far more common among Australian *Solanum* than previously thought and support the hypothesis that the combination of large fleshy fruits and endozoochorous dispersal represents a reversal in this study group. The general lack of direct evidence related to biotic dispersal (epizoochorous or endozoochorous) may be a function of declines and/or extinctions of vertebrate dispersers. Because of this, some taxa might now rely on secondary dispersal mechanisms (e.g. shakers, tumbleweeds, rafting) as a means to maintain current populations and establish new ones.

## Introduction

The large and cosmopolitan plant genus *Solanum* L. consists of nearly 1,400 accepted species [1], the majority of them exhibiting fleshy fruits linking them to biotic agents that disperse seeds as a consequence of frugivory [2]. Symon, [3] focusing only on the ca. 90 *Solanum* species described for Australia at the time, defined a set of fruit morphologies that correspond to hypothesized dispersal categories that can be summarized as follows: 1) pulpy/fleshy berries of various colors dispersed following ingestion (67 species), 2) firm, ultimately bony berries enclosed in a calyx with unclear dispersal (perhaps ingestion) (10 species), 3) smallish berries enclosed in a prickly calyx and ostensibly functioning as trample burrs on the feet of mammals (6 species), and 4) a postulated set of “oddball” fruits/mechanisms not matching those above (8 species).

A fair number of the species in categories 2, 3, and 4 belong to a recently-radiated [4,5] group of spiny solanums (*Solanum* subgenus *Leptostemonum*) known as the “*Solanum dioicum* group” [6,7], a set of species restricted to the Australian Monsoon Tropics (AMT) of northern Western Australia, the northern portions of the Northern Territory, and far-western Queensland. This species-group was first recognized by Whalen [7] as a diverse group of erect to spreading shrubs, and is the only group of *Solanum* species that includes both a large number of andromonoecious species and a set of cryptically dioecious species. Already considered unusual among Australian solanums for their breeding systems [8,9], these species are also markedly variable in morphology – with many taxa easily recognized via differences in vegetative form, habit, tomentum and armature, among other characters. Some of the greatest diversity, however, is related to fruit form and the degree to which fruits are enclosed by armed calyces, characters exhibiting particular influence on seed dispersal.

The 16 *S. dioicum* group taxa treated by Symon [3] include species that appear to disperse seeds via fruit fracturing (*S. oedipus*, *S. heteropodium*), censer-like mechanisms (*S. tudununggae*), or by catching in fur as trample burrs (*S. leopoldense*, *S. asymmetriphyllum*). The dispersal mechanisms of three other taxa were considered unclear by Symon (*S. carduiforme*, *S. petraeum*, *S. cataphractum*). The remaining species (9) were placed in the more typical fleshy-fruited category, particularly in the subcategory defined by Symon as having large (2-4 cm diameter), firm, yellowish berries at maturity (Table 1; Fig 1). Symon also assigned the trample burr character to members of the *S. echinatum* group (sensu Bean [8]), a clade of species known to be closely allied to the *S. dioicum* group based on more recent phylogenetic work [4,5].

**Table 1.**
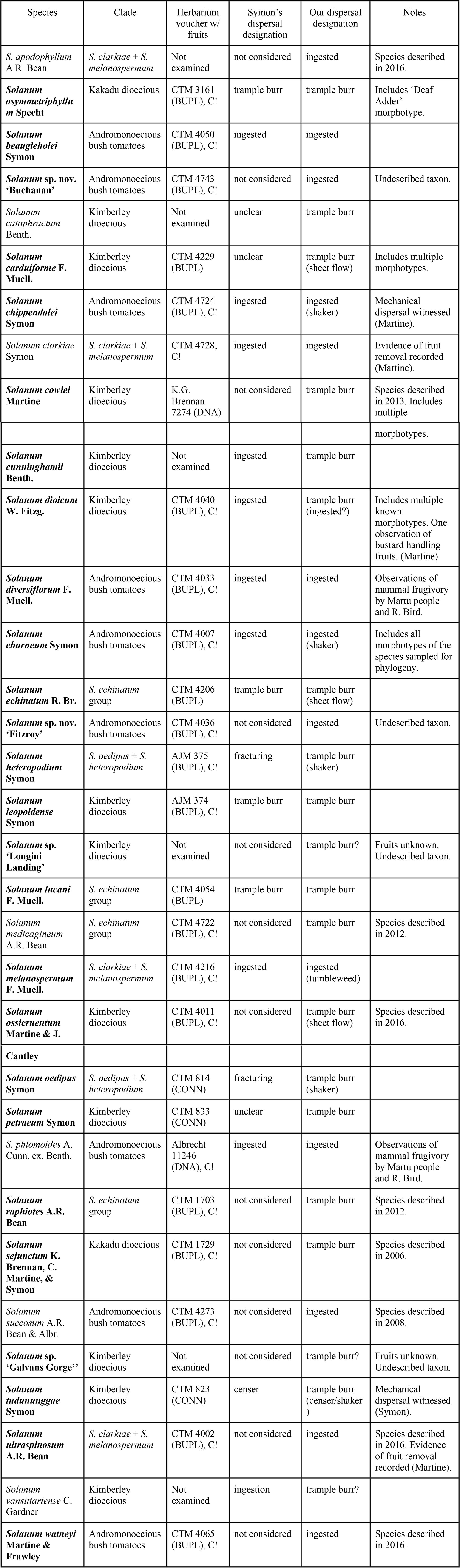
Species of the *S. dioicum* group and *S. echinatum* group sensu Bean [6] considered in this study.

**Figure 1.**
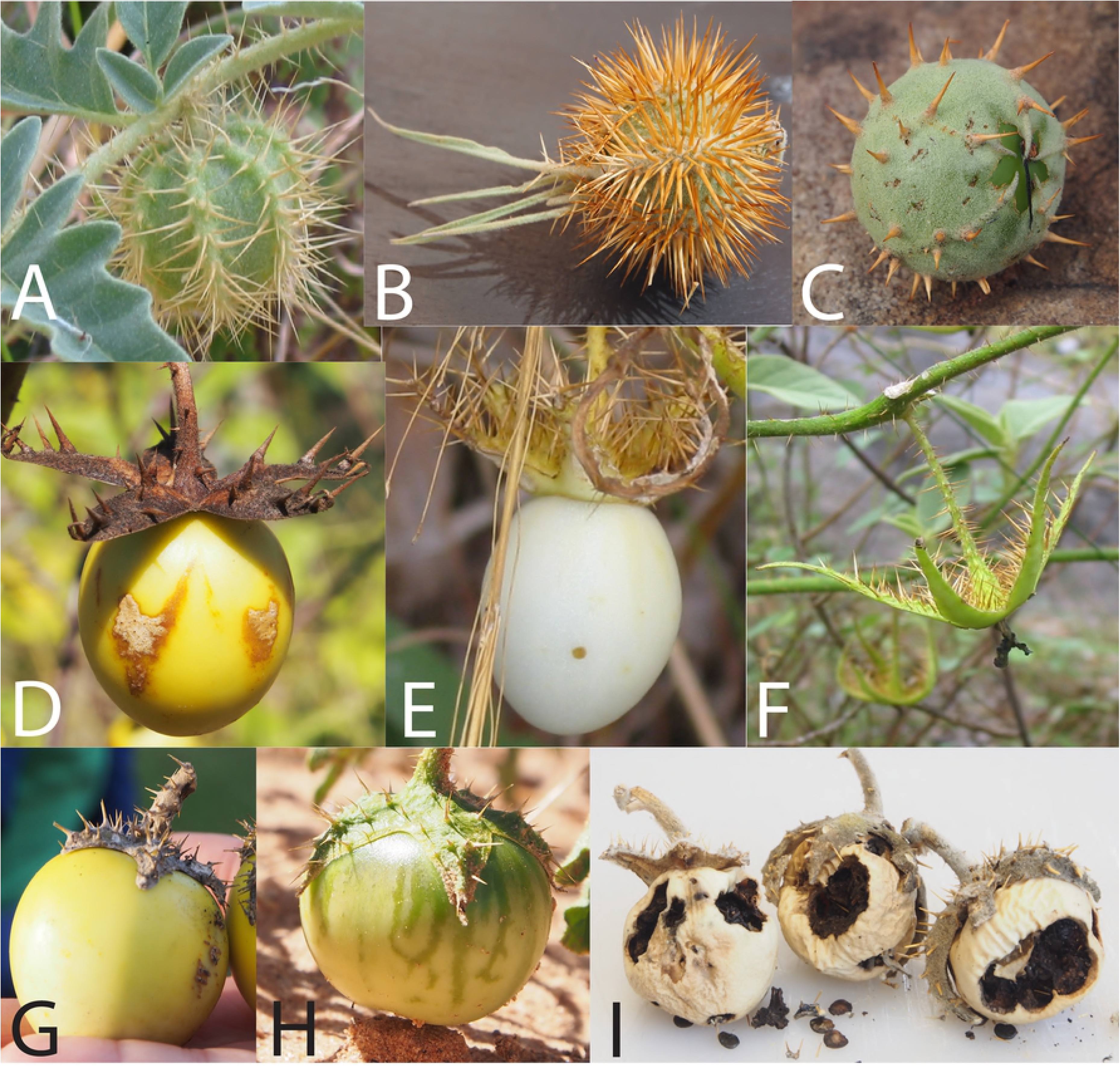
Fruit and calyx forms of selected AMT *Solanum* taxa. Photos A-C: Putative epizoochorous trample burr dispersal via accrescent prickly calyx (A: *S. carduiforme*, B: *S. ossicruentum*, C: *S. asymmetriphyllum*). Photos D-F: Putative ingestion dispersal after reflexing of accrescent calyx (D: *S. melanospermum*, E: *S. ultraspinosum* [fruit intact], F: *S. ultraspinosum* [fruit removed by unknown frugivore]. Photos G-I: Putative ingestion dispersal, calyx not enveloping fruit (G: *S. beaugleholei* [mature]; H: *S. diversiflorum* [immature, showing “cryptic coloration”], I: *S. chippendalei* [post-mature fruits exhibiting “shaker” mechanism]. Photos by C. Martine.

Associated fruiting herbarium vouchers (collector & number + herbarium acronym in parentheses; taxa marked with “C!” have also been cultivated at Bucknell as living specimens) along with dispersal methods as assigned by Symon [3] and the authors of this study (proposed secondary dispersal method in parentheses). Taxa included in the phylogenetic analyses are in bold. For taxa where fruiting specimens are unknown, dispersal designation is inferred based on phylogeny. (See S1 File for complete herbarium voucher and GenBank accession details.)

Calyx morphology is variable across the *S. dioicum* group, with calyces all prickly (sometimes heavily so) and accrescent to varying degrees of fruit coverage (illustrated in Fig 1). Fruits with calyces enveloping the fruit by half or less are assumed to fit into Symon’s broad category of ingested fruits, while those that are fully enveloped by calyces are assumed to be trample burrs. The exceptions to the latter condition recognized by Symon are 1) *S. tudununggae* and its censer mechanism, and 2) members of the *S. melanospermum* + *S. clarkiae* clade [5] in which the accrescent calyx reflexes and presents the berries at maturity – representing a sort of pre-dispersal defense mechanism to prevent the ingestion and distribution of immature seeds. Likewise, Symon [3] suggested that even immature fruits without enveloping calyces are cryptically colored (being green or striped green) and exhibit greater levels of alkaloids than those that are mature, a character present across the fleshy-fruited taxa (see Fig 1-H).

Specific seed dispersal mechanisms have not been published for any of the spiny *Solanum* species in the AMT. Seeds of some taxa can survive gut passage and are germinable after ingestion by rats ex situ [10], but throughout nearly 20 years of AMT *Solanum* field observations and inspection of thousands of wild plants by Martine and colleagues, frugivory (whether by direct observation or removed fruits) has rarely been witnessed (see Table 1). Indigenous knowledge of the biota of the AMT does, however, confirm that a few *Solanum* species are eaten by native mammals, especially macropods. The Martu people of the western desert report hill kangaroo (*Macropus robustus*) and burrowing bettong (*Bettongia leseuer,* now locally extinct) frugivory on *S. diversiflorum* and *S. phlomoides* [11] (R. Bird, pers comm). Local indigenous groups from the region of Kakadu National Park (a biodiversity hotspot in far-northern Northern Territory) suggest that the fruits of a few regional endemics may also occasionally be ingested by rock kangaroos [12], but only one species from the region’s flora (*Gardenia fucata*, Rubiaceae) has been found to be ingested and effectively (although rarely) dispersed by these reclusive marsupials [13]. Most AMT *Solanum* populations we have observed either retain the majority of their fruits well into the following season or, if the fruits are abscised when mature, the berries lay in uneaten piles beneath the parent plants. Given the limited extant evidence for frugivory/dispersal, one is left to ponder whether the production of large, fleshy berries by some AMT solanums represents an anachronism related to the mass extinction of nearly all large-bodied Australian animals over the last 400,000 years; or, perhaps, whether the production of such fruits is a relict of ancestral morphology and not related to biotic dispersal in Australia, at all.

Since Symon’s revision [14], many new species have been described from the region [15-22]. We here revisit seed dispersal strategies among the AMT *Solanum* taxa based on the present taxonomic understanding of the group and two decades of additional observations, primarily to test the hypothesis presented by Symon [3] that biotic dispersal by ingestion is the most common dispersal mechanism (with a few exceptions). Through new observations coupled with phylogenetic data we here show that not only are large, fleshy fruits less common than burr fruits in our study group, but also that they appear to represent a recent reversal in AMT *Solanum*.

## Methods

### Taxon sampling

Seventy-six individuals representing 50 species of *Solanum* were sampled from field collected dried tissue and/or fragments removed from pressed, dried herbarium specimens. This included 35 ingroup taxa to thoroughly sample clades and species-groups identified by Martine et al. [4,5] and to sample morphological diversity identified via fieldwork and specimen examination. We also sampled nine spiny *Solanum* outgroups, including three species from the closely-allied *S. echinatum* group sensu Bean [6] and six non-spiny *Solanum* outgroup species. Seven putative single-copy, intron-containing nuclear loci were newly identified and sequenced for all accessions (Jordon-Thaden, in prep; Table 2).

**Table 2:**
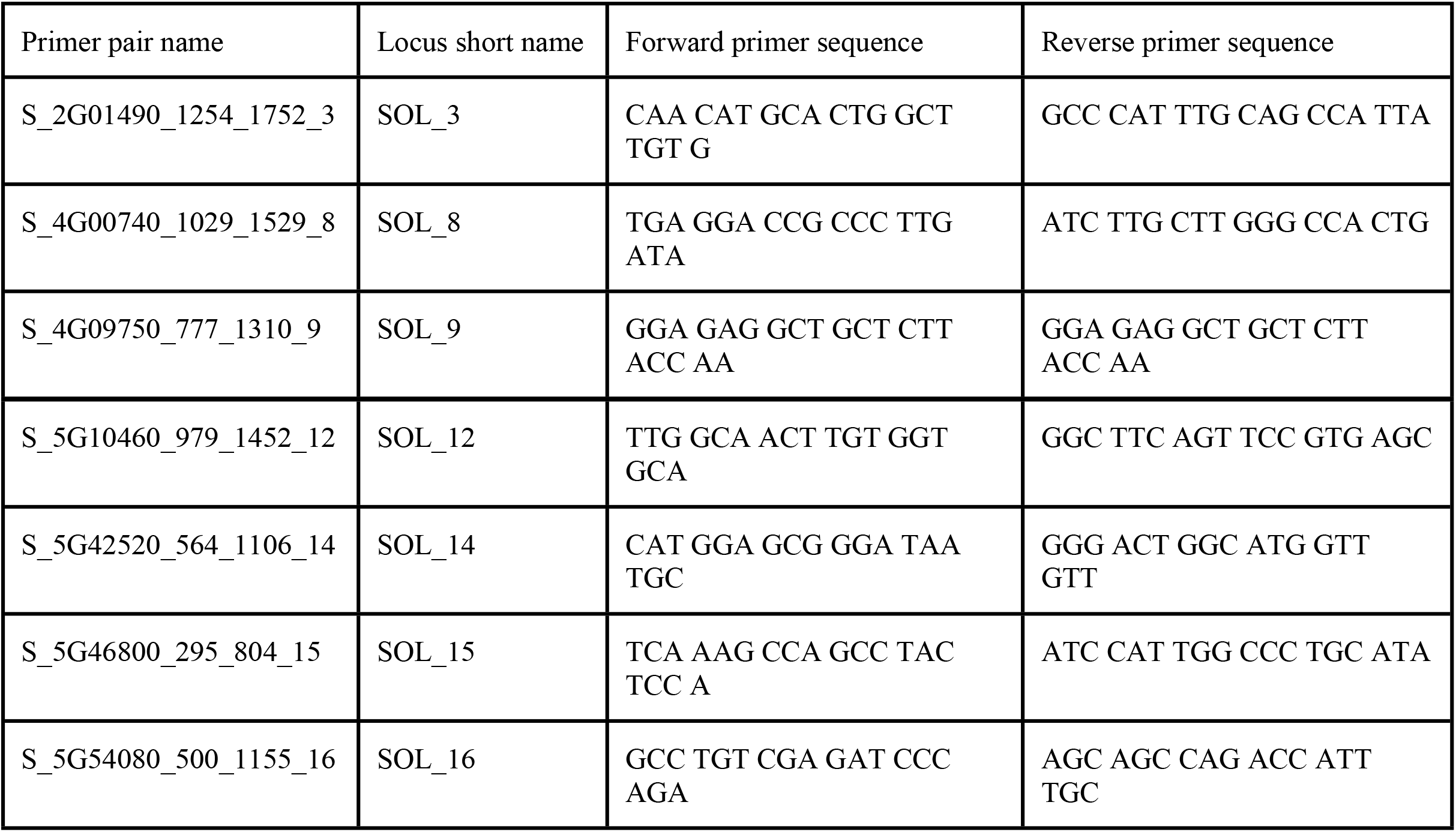
Primer sequences (5’-3’) used to amplify loci used in this study.

### Primer design, validation, & sequencing

Primers for this study were designed using MarkerMiner v.1.0 [23]. First, transcriptome assemblies for *Solanum dulcamara* L.*, S. ptycanthum* Dunal ex DC.*, S. cheesemanii* Geras.*, S. sisymbriifolium* Lam.*, S. lasiophyllum* Humb. & Bonpl. ex Dunal*, S. xanthocarpum* Schrad. & J.C. Wendl., were obtained from the 1 KP Project (www.onekp.org) and mapped to the *Arabidopsis thaliana* (L.) Heynh. genome. MarkerMiner v.1.0 was used to mine the resulting alignments for transcripts of single copy nuclear orthologs by filtering transcripts with scaffold length at least 900 bp and BLAST similarity at least 70% using both BLASTX and TBLASTN. The output was used to design primer pairs for the Fluidigm Access Array System (California, USA) using the Primer3 plugin [24] in Geneious (Biomatters Limited, New Zealand). Primers were selected that flanked predicted intronic regions with a size of 400-700 bp. Primer design and validation followed Uribe-Convers et al. [25] and the Access Array System protocol (Fluidigm, San Francisco, California, USA). Target-specific portions of primer pairs had a length of 20-25 bp, a melting temperature between 59 and 61° C, and contained no more than 3bp of homopolymer sequence for any nucleotide. Forty-eight primer pairs were selected for *Solanum*. To provide an annealing site for the Illumina sequencing adapters and sample-specific barcodes, a conserved sequence (CS) tag was added to the 5’ end of the forward and reverse primers at the time of oligonucleotide synthesis (CS1 for forward primers and CS2 for reverse primers [25]; purchased from Operon, Eurofins Scientific, Luxembourg).

To validate the primers, PCR amplification was conducted for 24 of the 48 primer pairs; validation reactions simulate the four-primer reaction of the Fluidigm microfluidic PCR system using a standard thermocycler. Reaction were carried out in 10 μl volumes each with final concentrations of reagents as follows: 1ng/μL gDNA (placed in master mix), 400 μM of each dNTP, 2x Fast Start High Fidelity reaction buffer (Roche), 9mM MgCl2 (Roche), 10% DMSO, 0.1U/μL FastStart High Fidelity Enzyme Blend (Roche), 2X AccessArray Loading Reagent (Fluidigm), 800nM AccessArray Barcode Primers for Illumina (designed by University of Idaho Institute for Bioinformatics and Evolutionary Studies Genomics Resources Core facility, iBEST), 200nM target specific primer mix (Operon), and water. PCR was conducted using following program: one round of 50°C for 2 minutes, 70°C for 20 minutes, 95°C for 10 minutes; ten rounds of 95°C for 15 seconds, 60°C for 30 seconds, 72°C for 1 minute; two rounds of 95°C for 15 seconds, 80°C for 30 seconds, 60°C for 30 seconds, and 72°C for 1 minute; eight rounds of 95°C for 15 seconds, 60°C for 30 seconds, 72°C for 1 minute; two rounds of 95°C for 15 seconds, 80°C for 30 seconds, 60°C for 30 seconds, 72°C for 1 minute; six rounds of 95°C for 15 seconds, 60°C for 30 seconds, 72°C for 1 minute; five rounds of 95°C for 15 seconds, 80°C for 30 seconds, 60°C for 30 seconds, 72°C for 1 minute. Each of the primer pairs were tested on three *Solanum* species (*S. asymmetriphyllum*, *S. eburneum*, and *S. elaeagnifolium*) and PCR products were visualized on 1.5% agarose gels run at 80V for 90 minutes. Validation was considered successful when, in all three taxa, amplification was observed as a single band that was not present in the negative and was within the appropriate size range [25].

Microfluidic PCR was carried out for 96 *Solanum* samples in the Fluidigm Access Array system (Fluidigm, San Francisco, California, USA) following the manufacturer’s protocols. Each of the 96 samples were amplified with the 24 primer pairs two times to completely fill the 48- well Fluidigm chip. To remove unused reagents and/or undetected primer dimers smaller than ~250 bp, each pool was purified with 0.6X AMPure XP beads (Agencourt, Beverly, Massachusetts, USA). PCR pools were analyzed in a Bioanalyzer High-Sensitivity Chip (Agilent Technologies, Santa Clara, California, USA) and standardized to 13 pM using the KAPA qPCR kit (KK4835; Kapa Biosystems, Woburn, Massachusetts, USA) on an ABI StepOnePlus Real-Time PCR System (Life Technologies, Grand Island, New York, USA). The resulting pools were multiplexed and sequenced in an Illumina MiSeq (San Diego, California, USA) to obtain 300 bp paired-end reads. Microfluidic PCR, downstream quality control and assurance, and sequencing were carried out at iBEST.

### Data processing

Raw reads were filtered, trimmed, and demultiplexed by barcode and target-specific primer using dbcAmplicons (https://github.com/msettles/dbcAmplicons [25]) and merged using FLASH [26]. Consensus sequences for each sample in all amplicons were generated using the reduce_amplicons R script (https://github.com/msettles/dbcAmplicons/blob/master/scripts/R/reduce_amplicons.R [25]).

### Phylogenetic Analyses

Individual loci were aligned using MUSCLE v3.8.5 [27] and adjusted manually. Phylogenetic topologies were estimated from the seven loci (summarized in Table 3) using three methods. First, gene trees for each locus were estimated using IQ-TREE version 1.5.5 [28,29] with the optimal substitution models selected by ModelFinder [30]. Clade support was determined by nonparametric bootstrapping using the ultrafast bootstrap with 1000 replicates [31]. Next, the resulting gene trees were used as input to ASTRAL-III [32] to estimate a species tree and multi-locus bootstrapping to calculate local posterior probability values for the species tree (100 replicates). Finally, we used Bayesian inference to estimate a phylogeny of a concatenated dataset using MrBayes 1.6.4 in parallel [33-35]. We applied the GTR+I+G model but allowed substitution rates for each gene to vary. The topology, branch lengths, shape, and state frequencies were unlinked. We sampled trees from two runs using eight chains (two hot, six cold) that ran for 5 × 108 generations with trees and model parameters sampled every 5000 generations. Convergence of the runs and stationarity were assessed for all parameters with the assistance of Tracer v.1.6.0 [36]. The Bayesian maximum clade credibility topology with posterior probabilities for each bipartition was summarized after discarding 25% of the sampled trees as burn-in using TreeAnnotator v.2.4.1 [37].

**Table 3:** Alignment characteristics by locus.

| Locus | # seqs in alignment | Ungapped length (bp) | Aligned length (bp) | Conserved sites | Variable sites | Parsimony informative sites | Missing data (%) |
| --- | --- | --- | --- | --- | --- | --- | --- |
| SOL_3 | 76 | 283-476 | 483 | 353 | 126 | 44 | 1.6 |
| SOL_8 | 80 | 458-465 | 466 | 313 | 130 | 47 | 1.1 |
| SOL_9 | 76 | 561 | 676 | 284 | 290 | 137 | 0 |
| SOL_1<br>2 | 84 | 267 | 267 | 161 | 106 | 65 | 0 |
| SOL_1<br>4 | 77 | 432-444 | 444 | 319 | 125 | 60 | 0 |
| SOL_1<br>5 | 64 | 565 | 673 | 377 | 228 | 120 | 0 |
| SOL_1<br>6 | 46 | 566 | 580 | 328 | 241 | 98 | 0 |

### Ancestral state reconstruction

We constructed an unordered morphological character matrix in Mesquite v.3.5 [38] for calyx form at fruit maturity, using the following character states: 1) reflexed, 2) subtended/typical, and 3) enclosed/trample burr. Character states were obtained for each taxon from Symon’s monograph [14] and/or field observations by the authors. We mapped calyx form onto the ASTRAL-III species tree estimate using unordered parsimony in Mesquite. The number and locations of transitions between character states was noted to determine whether or not calyx form is homoplasious and to better understand the associations between shifts in morphology within and among clades.

## Results

### Primer design and validation

MarkerMiner v.1.0 allowed for efficient design of primer pairs targeting putatively single-copy nuclear regions. We designed 48 pairs of primers for *Solanum*, of which 32 pairs were successfully validated and 24 were chosen for final amplification and sequencing in all samples (S1 File).

### Nuclear gene sequencing

Seven of the original 48 primer pairs we attempted obtained enough sequence data and coverage for downstream use (Table 2). In total, 503 new *Solanum* sequences were generated and analyzed. Data from this study are available in GenBank (S2 File).

### Phylogenetic analyses

Characteristics of the individual alignments used for phylogenetic analysis are summarized in Table 3. Individual gene trees estimated with IQ-TREE reveal weakly supported relationships among outgroup and ingroup clades and low overall resolution within sampled Australian *Solanum* (S1 Fig, trees available on Treebase xyz). No individual topologies lend support to the monophyly of the Australian species but do support some relationships within species and among close relatives. The relationships between many species and within species that are represented by multiple individuals are unresolved by the individual genes.

The ASTRAL-III species tree analysis was informed by the seven gene trees and attributed conflicts in the data to incomplete lineage sorting. This analysis uncovered a moderately well supported topology that is aligned with previous studies [4,5] (Fig 2). The ASTRAL-III topology shows Australian *Solanum* is divided into four lineages: the Kimberley dioecious clade, the Andromonoecious bush tomatoes, the Kakadu dioecious clade, and the *S. echinatum* group. The earliest diverging of these was the Kimberley dioecious clade (local posterior probability = 0.83). The Andromonoecious bush tomatoes were found to be polyphyletic in this study due to the position of *S. oedipus*, which occurred as a sister both to the Kakadu dioecious clade and to the rest of the Andromonoecious bush tomatoes. However, the remaining species of Andromonoecious bush tomatoes form a clade (local posterior probability <0.70). The Kakadu dioecious clade is well supported (local posterior probability = 0.94), and the *S. echinatum* group is moderately supported (local posterior probability = 0.79).

**Figure 2:**
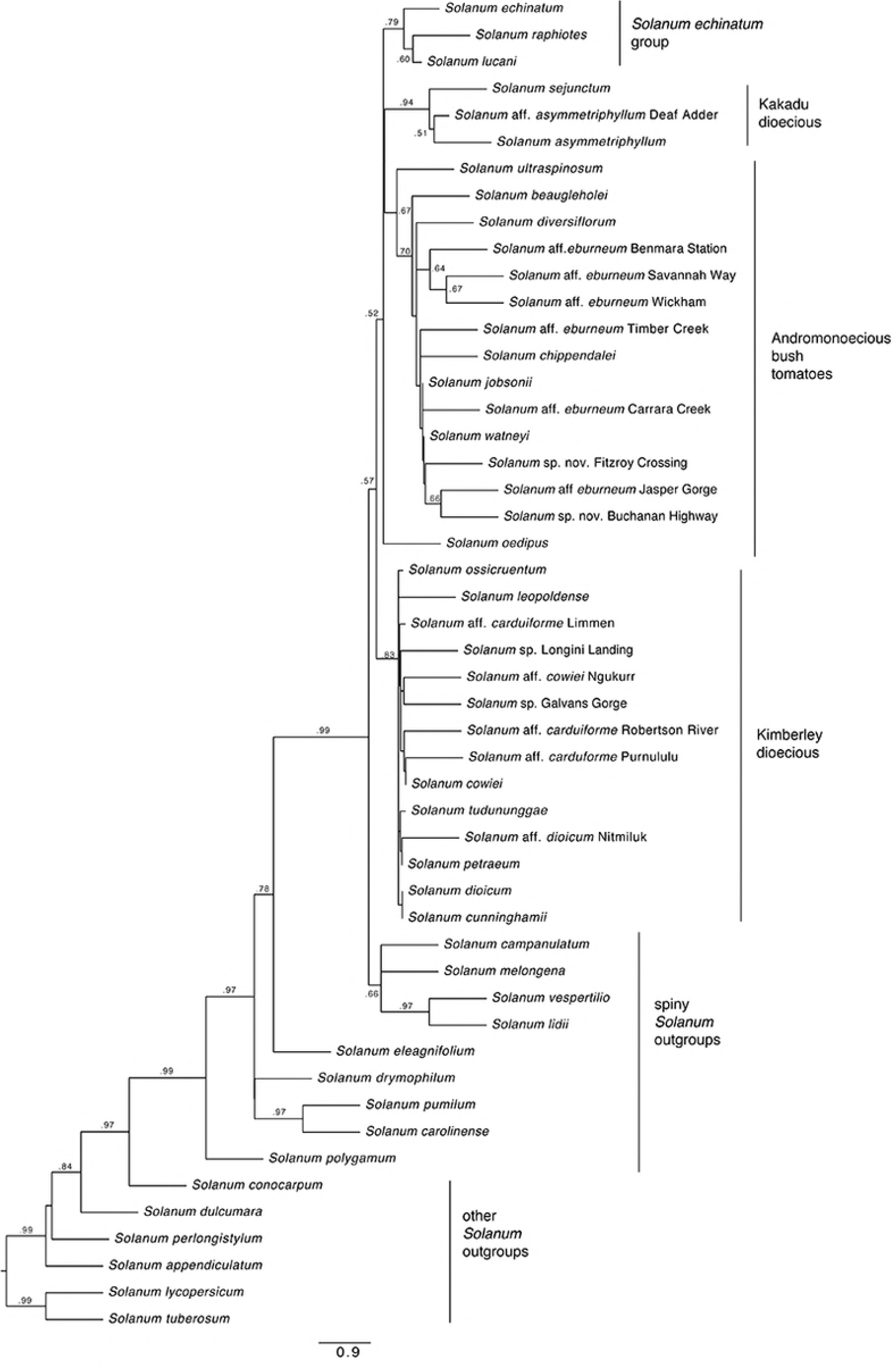
ASTRAL-III species tree generated from ML gene trees estimated in IQ-TREE. Values at nodes reflect local posterior probabilities of .50 or greater. Clade labels follow Martine, et al. (4,5). The *S. echinatum* group is identified sensu Bean [6].

An identical overall topology was recovered by the Bayesian maximum clade credibility tree (Fig 3) which was derived from a concatenated partitioned analysis of all loci, although the posterior probabilities (pp) from this analysis vary. The Kimberley dioecious clade was less well supported (pp < 0.70), while the clade of Andromonoecious bush tomatoes (excluding *S. oedipus*), the Kakadu dioecious clade, and the *S. echinatum* group were more well supported (pp of 0.99, 1.0, and 1.0 respectively). Also worth noting is the nonmonophyly of many species for which multiple accessions were included, a pattern also apparent in the individual gene trees (S1 Fig).

**Figure 3:**
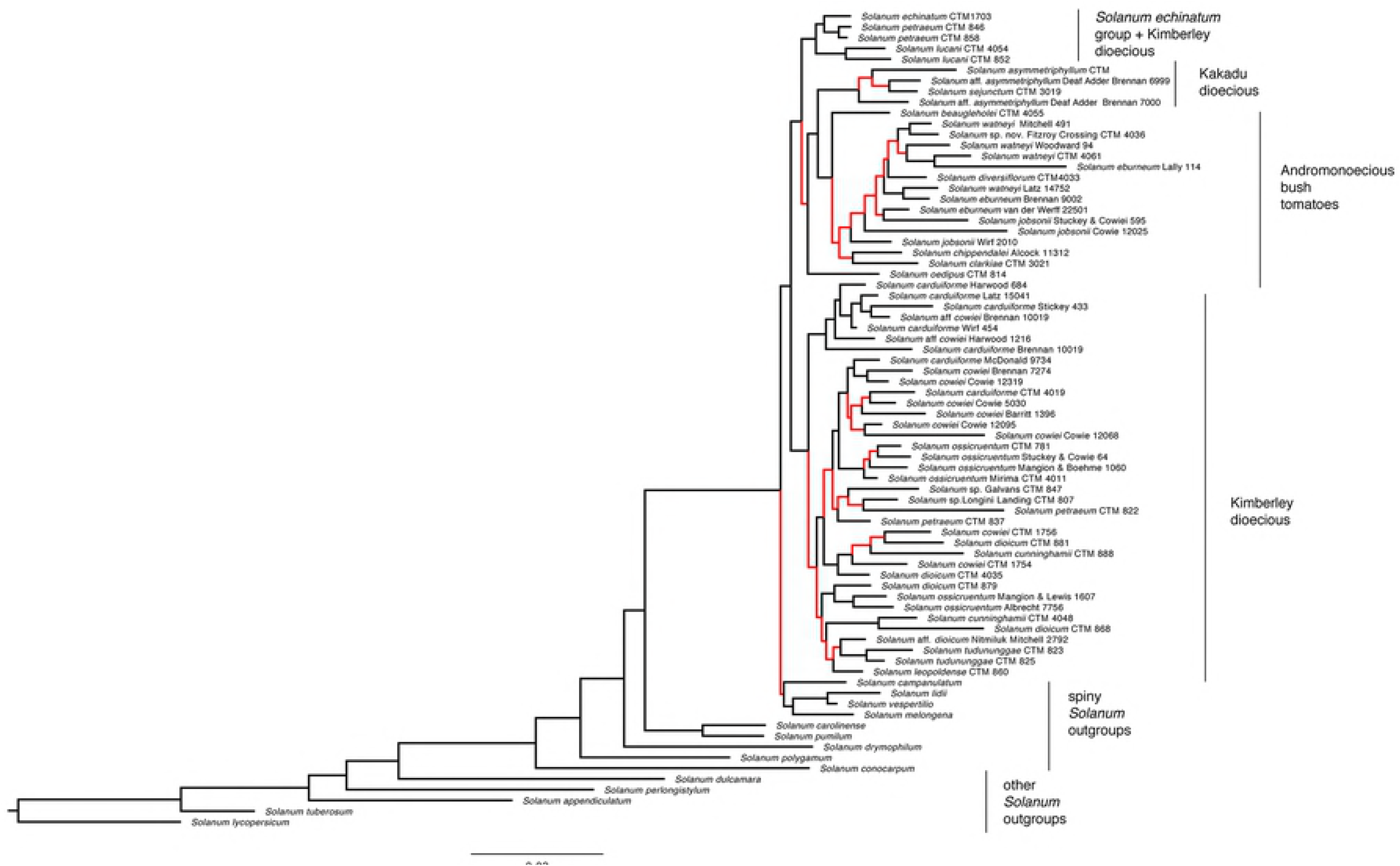
Maximum clade credibility topology inferred by Bayesian inference from concatenated partitioned loci. Red nodes reflect posterior probabilities of <0.9 and black nodes reflect posterior probabilities of 0.9-1. Clade labels follow Martine, et al. (4,5). The *S. echinatum* group is identified sensu Bean [6].

Overall, the phylogenetic analyses support a close relationship between the *S. dioicum* group sensu Whalen [7] and the *S. echinatum* group sensu Bean [6], and recover with some confidence three of the five clades within the *S. dioicum* group as identified by Martine et al. [5]: the Kimberley dioecious clade, the Kakadu dioecious clade, and the andromonoecious bush tomatoes. Of the two remaining clades not recovered here, the close relationship of the *S. melanospermum* + *S. clarkiae* clade to the bush tomatoes is inferred by the placement of the single species from the former clade in our dataset, *S. ultraspinosum*. The placement of *S. oedipus*, the single species included representing the *S. oedipus* + *S. heteropodium* clade, renders the bush tomatoes paraphyletic. Likewise, placement of the multiple morphotypes of *S. eburneum*, *S. carduiforme*, *S. dioicum*, and *S. cowiei* suggests that these species are not monophyletic as currently circumscribed and in need of revisionary study.

### Ancestral state reconstruction

The character state reconstruction (Fig 4) supports a typical or subtended calyx as ancestral in these *Solanum* species, including the spiny *Solanum* outgroup taxa. Within the Australian *S. dioicum* group, there is a transition to an enclosed or trample burr calyx morphology, which is the ancestral form in the *S. dioicum* group. There is equivocal support for a transition to reflexed calyx morphology or a reversal to the typical or subtended calyx morphology within most of the Andromonoecious bush tomatoes. *Solanum oedipus,* the andromonoecious species sister to the rest of the bush tomatoes plus the Kakadu dioecious taxa, retains the enclosed (trample burr) calyx form.

**Figure 4:**
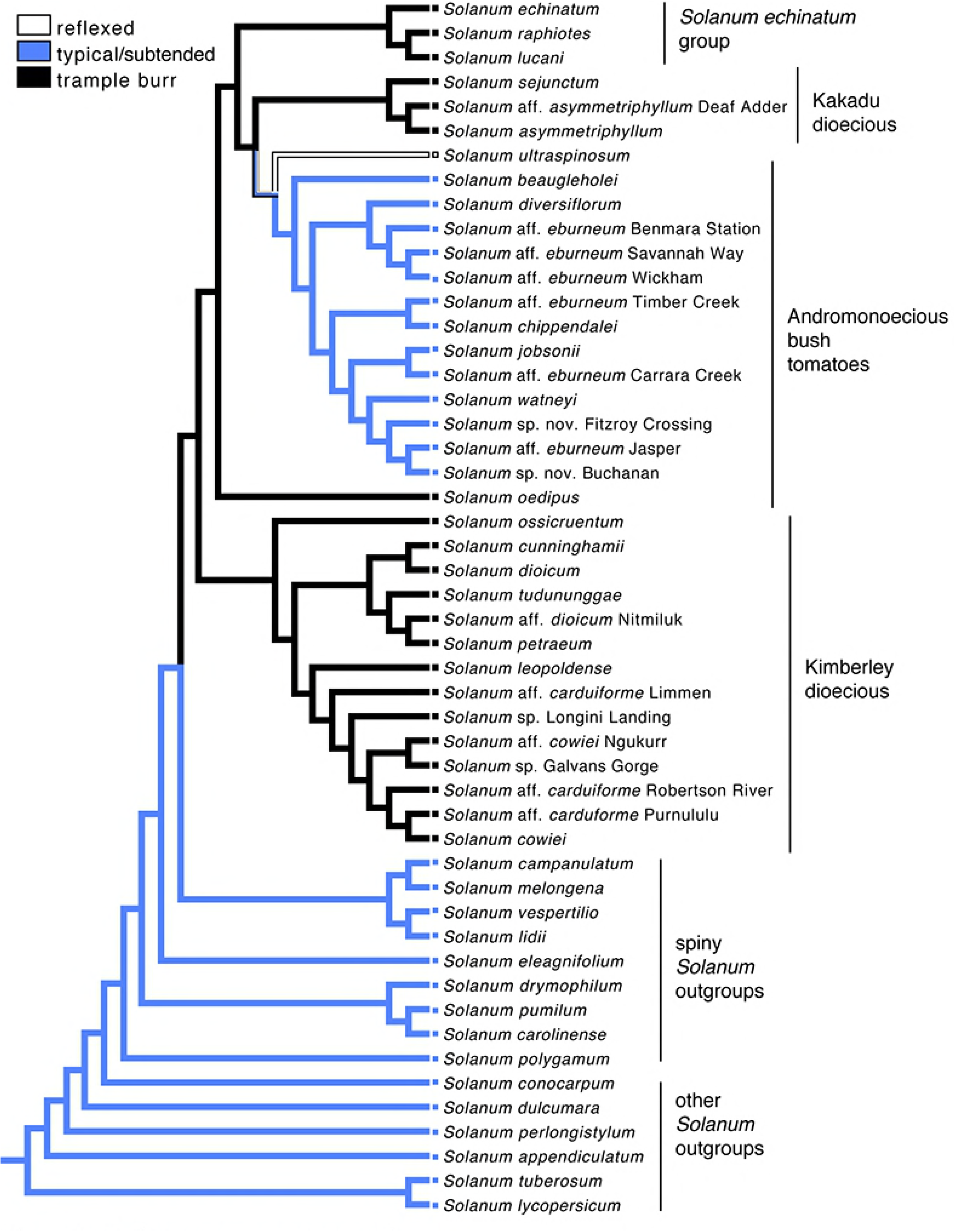
Most parsimonious ancestral state reconstruction of calyx morphology mapped onto the ASTRAL-III species tree. Clade labels follow Martine, et al. (4,5). The *S. echinatum* group is based on Bean [6].

## Discussion

We present a phylogeny of the *Solanum dioicum* group derived from nuclear data mined with MarkerMiner v.1.0 using 1KP transcriptomes with sequence data generated via microfluidic PCR-based amplicon enrichment and Illumina sequencing. This is the first study of the group to include intron-containing nuclear genes for phylogenetic analyses in the genus. These data support a monophyletic assemblage that includes the *S. dioicum* group species and reveals a close alliance with the *S. echinatum* group taxa. Future phylogenomic study of these plants will target members of both groups to test the close relationship uncovered here (McDonnell & Martine, in prep). The species tree hypothesis for these groups was used to investigate the evolution of calyx form within the lineage in an effort to better understand morphological diversity and natural history of these plants.

Calyx form and dispersal type are somewhat closely correlated with breeding systems among the species in what now might be considered the “*S. dioicum* + *S. echinatum* Group.” Species with large, fleshy fruits and mostly non-accrescent calyces are andromonoecious, while those with putative trample burr dispersal are dioecious or hermaphrodite. The phylogeny inferred by our analyses suggests that the andromonoecious breeding system (and thus their fruit type) is derived in our study group (having arisen from either dioecy or hermaphroditism), a hypothetical sequence that runs counter to the hermaphroditism-andromonoecy-dioecy sequence proposed by previous authors [3-5,8,9,14]. While this would upend conventional understanding of breeding system evolution in this group, we acknowledge that the limitations of our data force us to maintain skepticism pending additional study. The phylogeny presented here suffers from poor resolution which is a natural result from our use of a small number of short loci. We believe primer design challenges, such as the initial limitations of comparing *Solanum* transcripts to *Arabidopsis* genomes with MarkerMiner 1.0, could be a cause for the failure of many primer pairs to successfully amplify loci. However, we are confident that our current species-tree hypothesis for the group is plausible and that the combination of our phylogeny with many years of field observations offers worthwhile insight into the role of fruit and calyx form in seed dispersal among Australian *Solanum*.

### Fruit/calyx form and seed dispersal

Despite decades of study, the natural history of solanums in the AMT is still quite poorly understood – particularly with regard to plant-animal interactions including pollination [39] and seed dispersal [3]. Nearly all of the AMT *Solanum* species produce fleshy berries; and although traditional knowledge suggests that some animals do eat *Solanum* fruits [11,12] and may disperse seeds, no effective biotic seed dispersal interaction has yet been identified for a single species within the group. Still, one might infer that biotic dispersal following frugivory is a likely means by which many AMT solanums colonize new areas. Symon [3] suggested that birds, mammals, and lizards all play a role in endozoochorous dispersal of *Solanum* seeds in Australia; meanwhile, epizoochorous and non-biotic modes of dispersal (including wind, water, and mechanical modes) were hypothesized to play a secondary role.

As Symon pointed out, the interaction of fruit form and calyx morphology in seed dispersal appears to be significant. Based on a combination of field and greenhouse observations, herbarium study, experimental work, description of new species, and molecular phylogenetics over the last 15+ years, we suggest that for the *S. dioicum* group, where many species exhibit accrescent fruiting calyces for all or part of fruit/seed development, calyx form may be even more important than previously assumed. Specifically, current understanding of diversity in this group (including the many new species described since Symon’s 1981 monograph [14]) suggests that burr-fruited species (with accrescent calyces that remain closed after fruit maturity) are more common than calyces that only partly cover fleshy fruits. Whereas Symon hypothesized that six species across all of *Solanum* in Australia dispersed by trample burrs, we suggest that this number is closer to 18 in the *S. dioicum* group sensu stricto alone (Table 1). Likewise, the phylogenetic analyses infer that the *S. echinatum* group sensu Bean [6] (here represented by *S. echinatum*, *S. raphiotes*, and *S. lucani*) is nested in the *S. dioicum* group with which it shares the enclosed trample burr morphology (or, at least, an accrescent calyx). Recent work by Bean [40] suggests that species diversity in the *S. echinatum* group is greater than currently described and, therefore, so is the presence of trample burr dispersal in that clade.

For the species with accrescent calyces, the calyx appears to play two significant roles. One function, for most species, is in fruit and seed dispersal. The other function is likely that of a “pre-dispersal defense,” or the prevention of unripened fruits from being removed and ingested before the seeds are mature and germinable. In the case of some species, such as *S. ossicruentum*, readiness of seeds appears to correspond with whole fruits (with enveloping calyx) dropping off the plants entirely [21]. In perhaps the most elegant example of pre-dispersal defense, the members of the *S. melanospermum* + *S. clarkiae* Clade (those two species plus the recently-described *S. ultraspinosum* and *S. apodophyllum* [16]) enclose their unripened fruits in heavily-armed calyces before presenting them, still attached, with reflexed calyces at fruit maturity (Fig 1-E). This phenomenon is unusual in the family Solanaceae, where accrescent calyces are common but calyces reflexing to expose ripened berries is known from just two other genera [2] (*Jaltomata* and *Chamaesaracha*). Pre-dispersal defense via secondary chemistry among Australian species with non-accrescent calyces was proposed by Symon [3], with green striping on fruits produced by most of the non-accrescent species presumed to be both a warning signal and a camouflage via cryptic coloration against green foliage (Fig 1-H).

In addition to an assumed greater role for trample burr dispersal, our observations have also revealed that some of the secondary abiotic dispersal strategies proposed by Symon [3] are more common than previously imagined. Intriguingly, some of these seem to function as dispersal insurance for when fleshy fruits are not removed by frugivores and, instead, are left to dry out and eventually break apart. The mechanical “censer” dispersal Symon [3] ascribed to a single Australian species, *S. tudununggae,* might be better broadly described as a “shaker” mechanism inclusive of other species in which this has been observed. A prime example is *S. chippendalei*, where hundreds of individuals (near Taylor Creek, Northern Territory) were witnessed bearing the previous years’ fruits which, now dry and brittle, were dropping loose seeds from open fruit wall cavities (Fig 1-J) with the slightest touch or breeze (Martine, pers. obs.). Even in cultivation, *S. chippendalei* fruits age differentially – with some areas of the fruit wall breaking down much quicker than others as fruits pass beyond maturity. Similar shaker dispersal has also been seen in *S. eburneum* and may apply to what Symon^3^ described as “fracturing” in *S. oedipus* and *S. heteropodium*.

Symon [3] suggested that *S. pugiunculiferum*, a species found in tidal mudflats that is not a member of our study group, might be described as using tumbleweed dispersal – with whole plants (with fruits retained) breaking off and tumbling during the AMT dry season or rafting on water during the wet season. Our recent observations of *S. melanospermum* in the Northern Territory’s highly-dynamic Robinson and MacArthur River systems suggest that this species moves about in a similar fashion, even though it is also an obviously reflexed-calyx fruit-presenter (Fig 1-D). Shrubs holding onto hundreds of ripened and post-mature fruits have been found tossed along the deep sandy banks of these rivers like so much flotsam and jetsam. Water might likewise disperse some fruits that mature as bony and hollow, particularly when enclosed in calyces providing additional surface area among the prickles. We have observed the enclosed fruits of *S. ossicruentum*, *S. carduiforme*, and *S. echinatum* piled up in debris lines deposited by sheet wash from the previous season’s monsoon rains, a phenomenon not uncommon in arid zones [41-43].

Still, the broad observation that we have made is that, regardless of the “intended” mode of dispersal, precious few fruits appear to be carried off (whether internally or externally) by a biotic disperser and have thus become dependent on (seemingly) secondary abiotic forms of dispersal. Our work growing thousands of plants from seed in cultivation has shown that untreated seeds are able to germinate (even though rates are much higher following gibberellic acid treatments), suggesting that gut passage is not a dispersal requirement – even if it might offer some obvious benefits. Martine and Anderson [9] proposed that AMT *Solanum* might be engaged in a type of secondary dispersal by which macropods provide short-distance seed dispersal through a three-step process: fruit ingestion, “fecal seed storage” (after seeds are moved with animals to their denning sites and defecated), and “seasonal redispersal” via wet season rains. The general concept still applies to plant species producing trample burrs and may be even more likely than a scenario dependent on ingestion, given the hypothesized prevalence of epizoochory in the group.

The notion that the extinction of many of Australia’s medium- to large-bodied vertebrates in the last 400,000 years [44] may have rendered some primary dispersal mechanisms anachronistic is worth considering. One might imagine that the lack of current data related to effective seed dispersal is a simple reflection of the dearth of potential dispersers in the AMT, a factor made limiting by past megafaunal extinctions in Australia. However, no fossil record exists for large-scale megafauna extinctions over much of what is now the AMT [45,46]. Meanwhile, fossil records provide ample evidence for extinctions in southwestern [47] and southeastern [48] Australia; the assumed range of megafauna reached northward primarily along the western and eastern coasts [47,48], perhaps because of inconsistent availability of water in central and north-central regions.

In the areas of the continent where evidence is lacking for past megafauna abundance, relatively few extant plant species produce fruits fitting the typical profile of the megafauna dispersal system (large size, tough endocarp, and soft pulp [49]). Over much of interior and northern Australia, Pleistocene seed dispersal may have been left to macropods and other browsers (both extinct and extant), who may have been infrequently and opportunistically frugivorous – but who, largely, may have carried fruits/seeds in their fur. Trample burr fruit/calyx morphology may thus have been an advantage in an ecosystem lacking abundant large frugivores – and continues to be an advantage today.

Even where the megafauna syndrome is present in Australia it appears to not parallel the large-fruited big-tree version of Janzen and Martin [49]. Instead, plants exhibit a suite of growth forms that would have allowed shrubbier plants to resist browsing by large herbivorous animals – including physical and chemical defenses – while also producing fruits and seeds adapted for vertebrate dispersal [46], including trample burrs. In the present day, many of the remaining small- and medium-sized mammals (as well as frugivorous birds) throughout the AMT have recently experienced and continue to face declines [50-52]. This likely places many biotically-dispersed plants at risk and may put those species with potential secondary mechanisms (e.g. shakers or tumbleweeds) at a short-term advantage. In the long-run, however, local extirpations and wholesale extinctions of biotic dispersers are likely to have consequences for the plants linked to them. The observations of the Martu people [11] and R. Bird (pers.comm) at the boundary of the Little Sandy and Great Sandy Deserts are seemingly telling in this regard: one confirmed *Solanum* frugivore, the burrowing bettong (*Bettongia leseuer*), is now locally extinct; and the other, the likely-opportunistic hill kangaroo (*Macropus robustus*), handles the fruits of two species in such a way that the bitter seeds are avoided entirely. Demographic analyses (Cantley, et al., in prep) infer that several AMT *Solanum* species have undergone recent genetic bottlenecks, a trend that would likely continue should their biotic dispersers also go the way of the burrowing bettong.

The evolutionary development of calyx morphology is one of many exciting areas of current research in Solanaceae [53,54], especially in the context of seed dispersal biology. Among our *Solanum* study species, calyx form exhibits particular lability – with broad occurrence of derived accrescence/enclosure plus a reversal to the *Solanum* ancestral state of a non-enveloping calyx associated with fleshy berries. Further research employing larger molecular datasets (McDonnell and Martine, in prep) will shed greater light on the evolution of this and other fascinating elements associated with the reproductive biology of AMT *Solanum* species.

## Acknowledgements

This work would not have been possible without generous assistance from staff of the Northern Territory Herbarium, Palmerston (DNA) and the George Safford Torrey Herbarium, University of Connecticut (CONN), as well as assistance with collecting permits from Parks Australia, Northern Territory Parks and Wildlife Commission, and Western Australia Department of Parks and Wildlife. We gratefully acknowledge the Traditional Owners of the lands where many of our collections were made. T. Caton, M. Garanich, M. Spiro, W. Boop and others provided greenhouse support at Bucknell. Numerous individuals assisted with collection of specimens in the field during trips between 2004-2018, including R. Martine, I. Martine, J. Martine, E. Sullivan, D. Symon, P. Jobson, B. Barker, R. Barker, K. Brennan, E. Capaldi, D. Vogt, B. Figley, G. Lionheart, and B. Lavoie. Thank you to X reviewers for their valuable comments on earlier drafts of this paper.

## Supporting information

**S1 Fig. Individual gene trees for each of the seven loci as estimated by IQ-TREE.** Values at nodes reflect bootstrap support.

**S1 File. List of specimen vouchers and GenBank numbers for sequences used in this study.** For each accession, information is as follows: collector and collector number, locality, date of collection, acronym for herbarium (in parentheses) where voucher is held, and gene regions recovered for that accession. GenBank accession numbers for gene regions used in this study are listed for each taxon in the following order, with a dash (–) inserted where a region was not recovered: SOL3, SOL8, SOL9, SOL12, SOL14, SOL15, SOL16.

## References

1. Knapp S, Sagona E, Carbonell A, Chiarini F. A revision of the *Solanum elaeagnifolium* clade (Elaeagnifolium clade; subgenus Leptostemonum, Solanaceae). PhytoKeys. 2017;84: 1–104. doi:10.3897/phytokeys.84.12695

2. Hunziker AT. Genera Solanacearum: the genera of Solanaceae illustrated, arranged according to a new system. Königstein: Koeltz Scientific Books; 2001.

3. Symon DE. Fruit diversity and dispersal in *Solanum* in Australia. J Adel Bot Gard. 1979;1: 321–311. https://www.jstor.org/stable/23872220.

4. Martine CT, Anderson GJ, Les DH. Gender-bending aubergines: Molecular phylogenetics of cryptically dioecious *Solanum* in Australia. Aust Syst Bot. 2009;22: 107–120. doi: 10.1071/SB07039.

5. Martine CT, Vanderpool D, Anderson GJ, Les DH. Phylogenetic relationships of andromonoecious and dioecious Australian species of *Solanum* subgenus *Leptostemonum* section *Melongena*: inferences from ITS sequence data. Syst Bot. 2006;31: 410–420. https://www.jstor.org/stable/25064163.

6. Bean AR. The taxonomy and ecology of *Solanum* subg. *Leptostemonum* (Dunal) Bitter (Solanaceae) in Queensland and far north-eastern New South Wales, Australia. Austrobaileya. 2004;6: 639–816. http://www.jstor.org/stable/41739063.

7. Whalen MD Conspectus of species groups in Solanum subgenus Leptostemonum. Gentes Herbarum. 1984;12: 179–282.

8. Anderson GJ, Symon DE. Functional dioecy and andromonoecy in *Solanum*. Evolution. 1989;43: 204–219. doi: 10.1111/j.1558-5646.1989.tb04218.x.

9. Martine CT, Anderson GJ. Dioecy, pollination, and seed dispersal in Australian spiny *Solanum*. Acta Hortic. 2007;745: 269–283.

10. Lacey LM, Capaldi E, Jordon-Thaden I, Martine CT. Exploring the potential for *Solanum* fruit ingestion and seed dispersal by rock-dwelling mammals in the Australian Monsoon Tropics. Poster presented at: Botany Conference; 2015 Jul 25–29; Edmonton, Alberta. http://2015.botanyconference.org/engine/search/index.php?func=detail&aid=408.

11. Codding BF, Bliege Bird R, Kauhanen PG, Bird DW. Conservation or co-evolution? Intermediate levels of aboriginal burning and hunting have positive effects on kangaroo populations in Western Australia. Hum Ecol. 2014;42: 659–669. doi: 10.1007/s10745-014-9682-4.

12. Telfer WR, Garde MJ. Indigenous knowledge of rock kangaroo ecology in western Arnhem Land, Australia. Hum Ecol. 2006;34: 379–406. https://doi.org/10.1007/s10745-006-9023-3

13. Martine CT, Boni AJ, Capaldi EA, Lionheart G, Jordon-Thaden IE. Evidence of rock-dwelling macropod seed dispersal in a tropical monsoon community. North Territ Nat. 2016;27: 68–77.

14. Symon DE. A revision of genus *Solanum* in Australia. J Adel Bot Gard. 1981;4: 1–367.

15. Barrett RL. *Solanum zoeae* (Solanaceae), a new species of bush tomato from the North Kimberley, Western Australia. Nuytsia. 2013;23: 5–21.

16. Bean AR. Two new species of *Solanum* (Solanaceae) from the Northern Territory, Australia. Austrobaileya 2016;9: 524–533.

17. Bean AR, Albrecht DE. *Solanum succosum* A.R.Bean & Albr. (Solanaceae), a new species allied to *S. chippendalei* Symon. Austrobaileya. 2008;7: 669–675. http://www.jstor.org/stable/41739087.

18. Brennan K, Martine CT, Symon DE. *Solanum sejunctum* (Solanaceae), a new functionally dioecious species from Kakadu National Park, Northern Territory, Australia. The Beagle, Records of the Museums and Art Galleries of the Northern Territory. 2006;22: 1–7.

19. Lacey LM, Cantley JT, Martine CT. *Solanum jobsonii*, a novel andromonoecious bush tomato from a new Australian national park. PhytoKeys. 2017;82: 1–13. doi: 10.3897/phytokeys.82.12106

20. Martine CT, Frawley ES, Cantley JC, Jordon-Thaden IE. *Solanum watneyi*, a new bush tomato species from the Northern Territory, Australia named for Mark Watney of the book and film “The Martian”. PhytoKeys. 2016a;61: 1–13. doi: 10.3897/phytokeys.61.6995

21. Martine CT, Cantley JC, Frawley ES, Butler A, Jordon-Thaden IE. New functionally dioecious bush tomato from northwestern Australia, *Solanum ossicruentum*, may utilize “trample burr” dispersal. PhytoKeys. 2016b;63: 19–29. doi: 10.3897/phytokeys.637743

22. Martine CT, Symon DE, Capaldi Evans E. A new cryptically dioecious species of bush tomato (*Solanum*) from the Northern Territory, Australia. PhytoKeys. 2013;30: 23–31. doi: 10.3897/phytokeys.30.6003

23. Chamala S, García N, Godden GT, Krishnakumar V, Jordon-Thaden IE, De Smet R, et al. MarkerMiner 1.0: A new application for phylogenetic marker development using angiosperm transcriptomes. Applications in Plant Sciences, 2015;3(4): 1400115. doi:10.3732/apps.1400115

24. Untergasser A, Cutcutache I, Koressaa T, Ye J, Faircloth BC, et al. Primer3—new capabilities and interfaces. Nucleaic Acids Res. 2012;40(15): e115. doi: 10.1093/nar/gks596.

25. Uribe-Convers S, Settles ML, Tank DC. A Phylogenomic Approach Based on PCR Target Enrichment and High Throughput Sequencing: Resolving the Diversity within the South American Species of *Bartsia* L. (Orobanchaceae). PLoS ONE. 2016;11(2): e0148203. doi:10.1371/journal.pone.0148203

26. Magoc T, Salzberg S. FLASH: Fast length adjustment of short reads to improve genome assemblies. Bioinformatics. 2011;27: 2957–63. doi: 10.1093/bioinformatics/btr507.

27. Edgar R. 2004. MUSCLE: multiple sequence alignment with high accuracy and high throughput. Nucleic Acids Research. 2004;32: 1792–1797. doi: 10.1093/nar/gkh340.

28. Chernomor O, von Haeseler A, Minh BQ. Terrace aware data structure for phylogenomic inference from supermatrices. Syst Biol. 2016;65: 997–1008. doi: 10.1093/sysbio/syw037.

29. Nguyen LT, Schmidt HA, von Haeseler A, Minh BQ. IQ-TREE: A fast and effective stochastic algorithm for estimating maximum-likelihood phylogenies. Mol Biol Evol. 2015;32(1): 268–274. doi:10.1093/molbev/msu300

30. Kalyaanamoorthy S, Minh BQ, Wong TKF, von Haeseler A, Jermiin LS. ModelFinder: fast model selection for accurate phylogenetic estimates. Nat Methods. 2017;14: 587–589. doi: 10.1038/nmeth.4285.

31. Minh BQ, Nguyen MAT, von Haeseler A. Ultrafast approximation for phylogenetic bootstrap. Mol Biol Evol. 2013;30: 1188–1195. doi: 10.1093/molbev/mst024.

32. Zhang C, Rabiee M, Sayyari E, Mirarab S. ASTRAL-III: polynomial time species tree reconstruction from partially resolved gene trees. BMC Bioinformatics. 2018;19(6): 153. doi:10.1186/s12859-018-2129-y

33. Altekar G, Dwarkadas A, Huelsenbeck JP, Ronquist F. Parallel Metropolis-coupled Markov chain Monte Carlo for Bayesian phylogenetic inference. Bioinformatics. 2004;20: 407–415. doi: 10.1093/bioinformatics/btg427.

34. Huelsenbeck JP, Ronquist F. MRBAYES: Bayesian inference of phylogeny. Bioinformatics. 2001;17: 754–755.

35. Ronquist F, Huelsenbeck JP. MRBAYES 3: Bayesian phylogenetic inference under mixed models. Bioinformatics. 2003;19: 1572–1574.

36. Rambaut A, Suchard MA, Xie D, Drummond AJ. Tracer. Version 1.6. 2014. Available from: http://tree.bio.ed.ac.uk/software/tracer.

37. Rambaut A, Drummond AJ. TreeAnnotator. Version 2.4.1. 2016. Available from: http://beast.bio.ed.ac.uk/treeannotator.

38. Maddison WP, Maddison DR. Mesquite: a modular system for evolutionary analysis. Version 3.51. 2018. Available from: http://www.mesquiteproject.org.

39. Anderson GJ, Symon DE. Insect foragers on *Solanum* flowers in Australia. Ann Mo Bot Gard. 1988; 75: 842–852. doi: 10.2307/2409175

40. Bean AR. A taxonomic revision of the *Solanum echinatum* group (Solanaceae). Phytotaxa. 2012;57: 33–50. doi:10.11646/phytotaxa.57.1.6.

41. Mott JJ, McComb AJ. Patterns in annual vegetation and microrelief in an arid region of Western Australia. J Ecol. 1974;62: 115–26. doi: 10.2307/2258884.

42. Reichman JO. Spatial and Temporal Variation of Seed Distributions in Sonoran Desert Soils. J Biogeogr. 1984;11: 1–11. doi:10.2307/2844771

43. Venable DL, Flores-Martinez A, Muller-Landau HC, Barron-Gafford G, Becerra JX. Seed dispersal of desert annuals. Ecology. 2008;89(8): 2218–2227. https://www.jstor.org/stable/27650747.

44. Wroe S, Field JH, Archer M, Grayson DK, Price GJ, Louys J, et al. Climate change frames debate over the extinction of megafauna in Sahul (Pleistocene Australia - New Guinea). PNAS; 110: 8777–8781. doi: 10.1073/pnas.1302698110.

45. Horton DR. Red kangaroos: last of the Australian megafauna. In: Martin PS, Klein RG, eds. Quaternary extinctions: a prehistoric revolution. Tucson: University of Arizona Press; 1984, pp. 639–690.

46. Johnson C. Australia’s mammal extinctions: a 50,000 year history. Cambridge: Cambridge University Press; 2006.

47. Merilees D. Comings and goings of late Quaternary mammals in extreme southwestern Australia. In: Martin PS, Klein RG, eds. Quaternary extinctions: a prehistoric revolution. Tucson: University of Arizona Press; 1984, pp 629–638.

48. Murray P. Extinctions downunder: a bestiary of extinct Australian late Pleistocene monotremes and marsupials. In: Martin PS, Klein RG, eds. Quaternary extinctions: a prehistoric revolution. Tucson: University of Arizona Press; 1984.

49. Janzen DH, Martin PS. Neotropical anachronisms: the fruits the gomphotheres ate. Science. 1982;215: 19–27. doi: 10.1126/science.215.4528.19.

50. Woinarski J, Milne DJ, Wanganeen G. Changes in mammal populations in relatively intact landscapes of Kakadu National Park, Northern Territory, Australia. Austral Ecol. 2001;26: 360–370. doi: 10.1046/j.1442-9993.2001.01121.x.

51. Woinarski J, Fisher A. Threatened terrestrial animals of Kakadu National Park: Which species?; How are they faring? And what needs to be done for them? In: Winderlich S, Woinarski J, eds. Kakadu National Park Landscape Symposia Series. Symposium 7: Conservation of threatened species. Internal Report 623. Darwin: Supervising Scientist; 2013;93–104.

52. Ziembicki MR, Woinarski JCZ, Mackey B. Evaluating the status of species using Indigenous knowledge: Novel evidence for major native mammal declines in northern Australia. Biol Conserv. 2013;157: 78–92. doi: 10.1016/j.biocon.2012.07.004.

53. Deanna R, Larter MD, Barboza GE, Smith SD. Repeated evolution of a morphological novelty: a phylogenetic analysis of inflated fruiting calyx in the Physalidae tribe (Solanaceae). 2018; bioRxiv 425991; doi: 10.1101/425991.

54. Knapp S. Tobacco to tomatoes: a phylogenetic perspective on fruit diversity in the Solanaceae. J. Exp. Bot. 2002;53: 2001–2022. doi: 10.1093/jxb/erf068.

